# Antimicrobial Resistance Profiles of Bacterial Contaminants from a Tertiary Hospital in Kenya: An Urgent Call for Action Against the Global Threat of Antimicrobial Resistance

**DOI:** 10.1101/2023.05.11.540441

**Authors:** Kolek Chester, Kavulavu Briton, Faith Okalebo, Benson Singa, Mary Masheti, Ian Omuom, Ochieng Odhoch, Chris Oduol, Robert Musyimi, Caroline Tigoi, Kirkby D Tickell

## Abstract

**Background:** Hospital-acquired infections (HAIs) represent the most prevalent adverse event among patients in hospital settings. Contamination with pathogenic bacteria that are highly resistant in the hospital environment increases the risk of HAIs.

**Objective:** The antimicrobial resistance (AMR) patterns of hospital contaminants isolated from highly frequented surfaces in a tertiary hospital in Kenya.

**Methods:** A total of 62 swabs were collected from selected surfaces, equipment, and health workers’ palms in April 2020. They were cultured and bacterial contaminants were identified using standard microbiological procedures and their AMR patterns were determined using recommended laboratory assays.

**Results:** Of the 62 swabs collected, 61.3% (n=38) yielded bacterial growth, from which 46 bacteria were isolated. Swab positivity varied across the departments as follows: gynecology wards (78.6%), New Born Unit (NBU) (56.2%), Pediatric ward (61.9%), and Renal Unit (45.5%). Gram negative species comprised 86.96%(n=40) while Gram positive species comprised 13.04%(n=6). Of all the 46 isolates obtained, 36.96% (n=17) were positive for the resistance markers screened. Specifically, 10.9% (n=5) showed both extended-spectrum beta-lactamases (ESBL) and carbapenem-resistant (CR) resistance, while 23.9%(n=11) were positive for ESBL production. The rest were non-resistant strains as shown by negative ESBL at 47.8% (n=22), methicillin sensitivity at 13% (n=6) and vancomycin sensitivity at 2.2% (n=1). *Acinetobacter* species which were most reported, had the highest resistance (36.84% (7/19).

**Conclusion:** There was a high prevalence of contamination with resistant pathogenic bacteria species. *Acinetobacter* species were the most common pathogen. Interventions are needed to mitigate the problem of resistant HAI.

## INTRODUCTION

Hospital-acquired infections (HAIs) continue to threaten the treatment outcomes of many patients worldwide. According to World Health Organization (WHO), HAIs represent the most prevalent adverse event among patients in hospital settings (1). Morbidity and mortality, high costs of care and reduced quality of life, are among the detrimental effects of HAIs (2–4). Over million cases of HAIs are reported annually in Europe (2). These cases often result in deaths.

About 90,000 deaths are to be attributable to the six most common types of HAIs: pneumonia; urinary tract infection (HA-UTI); surgical site infections (SSI); healthcare-associated *Clostridium difficile*; neonatal sepsis; and bloodstream infection(2). Whereas the average global prevalence of HAIs is about 0.14%(3), the prevalence is even higher in Africa and the developing world. According to a meta-analysis by Abubakar et al.(3), the pooled prevalence of HAIs in Africa was 12.76%.

The continued isolation of multidrug-resistant pathogens among HAI cases is an alarming finding. In one study that assessed the antimicrobial susceptibility (AST) patterns among HAI cases in China, Zhang et al. (5). observed an HAI prevalence of 12.4%, of which 0.18% were multidrug-resistant (MDR) pathogens. The most common resistant pathogens were *Acinetobacter spp* (53.86%) and *Pseudomonas spp*. (21.6%) (5). In a global study by Murray et al. (6), it was observed that the reporting of resistant pathogens was high. Of these resistant pathogens, *Escherichia coli*, coagulase-negative staphylococci (CONS), Staphylococcus *spp*, and Pseudomonas species were the most widely reported (6).

The ward in which a patient is a possible risk factor for HAIs. According to Despotovic et al. (7), up to 32.7% of patients in an intensive care unit had HAIs of which over 50% were caused by organisms resistant to common antibiotics. Local data appears to point to a similar picture. Kagia et al. (7) observed a carriage rate of 10% for extended-spectrum beta-lactamase (ESBL) producing pathogens among neonates in a hospital in Kilifi. Among the positive ESBL cases, 55% had acquired ESBL pathogens in the hospital (8). Among surface contaminants, the prevalence of MDR pathogens is not different. According to a study that assessed surface contamination in multiple hospitals in Kenya, Odoyo et al.(9) reported that up to 12.6% of the sampled high-touch surfaces were contaminated with multidrug resistant (MDR) pathogens, the most common of which was *Acinetobacter boumanii* at 3.7%. Others were *Klebsiella. pneumoniae* (3.6%), *Enterobacter* species (3.1%), methicillin-resistant *Staphylococcus. aureus* (MRSA) (0.8%), *Escherica coli* (0.8%), *Psuedomonas aeruginosa* (0.3%), and *Enterococcus faecalis* and *E. faecium* (0.3%). It is to be noted that these bacteria are members of the highly stubborn ESKAPE group.

Much of the literature available on the AMR patterns of HAIs considered patient isolates mainly. Odoyo et al. (8) reported that surface contamination with resistant pathogens increases the risk for difficult-to-treat HAIs among hospitalized patients. Thus, any hospital intending to implement accurate and effective Infection Prevention and Control (IPC) strategies must understand the prevalence and AMR patterns of surface contaminants in the highly-frequented surfaces such as sinks, door handles, staff hands, and commonly used and shared equipment (9). These surfaces have been reported as significant agents in infection spread in the hospital setting and are hence are the focus of hospital IPC guidelines(10–12). The present study aimed to determine the antimicrobial susceptibility patterns of surface contaminants at the Migori County Referral Hospital, Kenya.

## METHODS

A descriptive cross-sectional study was conducted at Migori County Referral Hospital (MCRH) pediatric and gynecology wards and the Renal and Newborn Units. The facility is a 150-bed capacity hospital in Migori County located southwest of Kenya (1.06412^0^S,34.47573^0^E) at the Kenya-Tanzania border. It is the highest referral hospital in Migori County.

Purposive sampling was used to select highly frequented sites that would pose an increased threat for infection spread, including door handles, healthcare workers’ hands, sinks, and shared equipment. Whereas the sampling approach targeted a minimum sample size of 50 (20 from equipment, 20 from hospital surfaces, and ten from staff), 62 samples were collected, as detailed in **Table 1**.

**Table 1.** Departments and types of surfaces from which of swabs were obtained

| Type of Specimen | Department |  |  |  | Total |
| --- | --- | --- | --- | --- | --- |
|  | Gynecology | New Born Unit | Pediatric | Renal |  |
| Equipment | 5 | 8 | 7 | 5 | 25 |
| Staff | 4 | 4 | 7 | 4 | 19 |
| Surface | 5 | 4 | 7 | 2 | 18 |
| Total | 14 | 16 | 21 | 11 | 62 |

Surfaces were swabbed with moistened (sterile 8.5% normal saline) COPAN floqswabs®. The swab was spread over the surface to cover about 30 cm^2^ touches. After collection, the swabs were transported in Cary-Blair medium tube at 2-8°C in a cool box packed with ice cubes. Upon arrival in the laboratory, tubes were vortexed at 300 RPM for 20 seconds and opened inside Class II biosafety cabinets to reduce exposure to aerosols. Twenty microliters of broth was inoculated and streaked on horse blood agar and cystine–lactose–electrolyte-deficient (CLED) agar plates. Horse blood agar and CLED agar plates were incubated at 37°C in 5% CO_2_ and aerobic incubator for 24 hours. Any emerging colonial growth was sub-cultured on respective plates and set as mentioned above.

Identification of bacterial genus and species were done by MALDI-TOF at the KEMRI Wellcome Trust Laboratory in Kilifi, and antibiotic susceptibility testing was done on all pathogens by disc diffusion (Kirby Bauer) as described in Clinical and Laboratory Science Institute (CLSI) guidelines to identify antibiotic-associated resistant markers. ESBL production was screened phenotypically using the double-disk synergy method. Cefotaxime (30 mcg) and ceftazidime (30 mcg) were used alone for screening before double confirmation using cefotaxime/ clavulanate (30/10 mcg) and ceftazidime/clavulanate (30/10 mcg) combination as per the recommendation of CLSI guidelines (Figure 1). Gram negative species were subjected to ESBL, CRO, and Vancomycin Resistance screening whereas the gram positive species were subjected to Methicillin Resistance screening according to conventional procedures.

**Figure 1.**
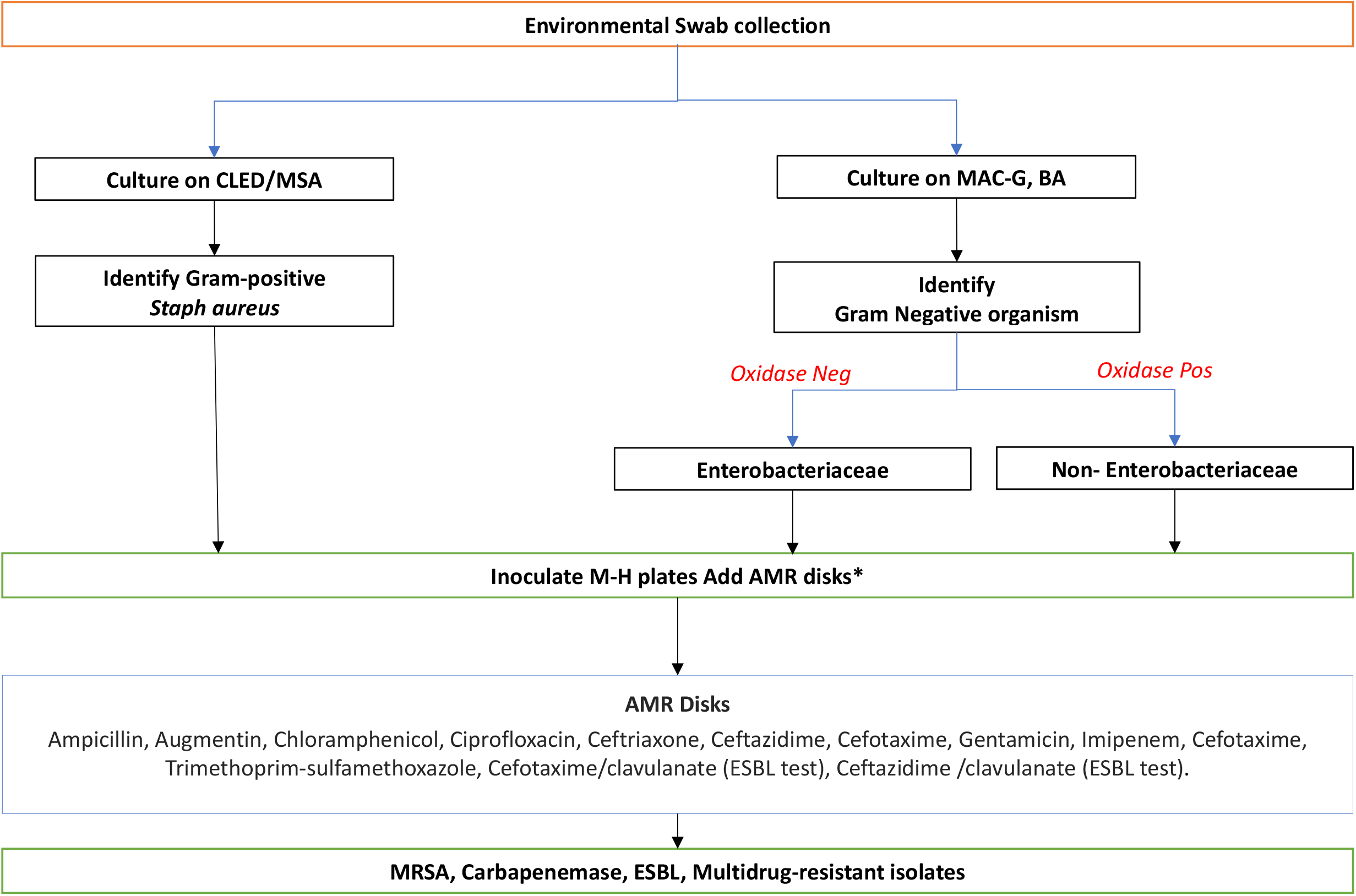
Flow chart summarizing the procedure followed for isolation, identification, and antibiotic susceptibility characterization of the specimen collected from MCRH

## RESULTS

Out of the 62 swabs collected, 61.3%(38 swabs) showed positive growth, as summarized in **Table 2**. The gynecology ward had the highest prevalence of contaminated swabs followed by the pediatric ward.

**Table 2.** Prevalence of swabs that has microbial contamination in each department

| Department | Total Swabs | Positive Swabs | % Positivity |
| --- | --- | --- | --- |
| Gynecology | 14 | 11 | 78.6 |
| New Born Unit | 16 | 9 | 56.2 |
| Pediatric | 21 | 13 | 61.9 |
| Renal Unit | 11 | 5 | 45.5 |
| **Total** | **62** | **38** | **61.3** |

A total of 46 isolates were obtained. Of the 46 isolates obtained, a majority were *Acinetobacter spp* which comprised 41.3% (n=19) (**Table 3**). Regarding AMR patterns, 36.96% (n=17) were positive for the resistance markers screened. Important to note as the fact that among the *Acinetobacter spp*, the prevalence of resistance rate 36.8% (7/19). Gram negative species comprised 87.0%(n=40) while Gram positive species comprised 13.0%(n=6). Among the majority of gram negative species, the prevelance of resistance was 42.5% (17/40). Worth notice is the fact that none of the gram-positive bacteria were resistant. (**Table 4**).

**Table 3.** Summary of the resistance patterns of the isolates obtained (by Genus).

| **Genus** | **Total** | **%** | **Resistant** | **%** |
| --- | --- | --- | --- | --- |
| *Acinetobacter* | 19 | 41.3 | 7 | 36.84 |
| *Staphylococcus* | 6 | 13.04 | 0 | 0.00 |
| *Enterobacter* | 5 | 10.9 | 4 | 80.00 |
| *Citrobacter* | 1 | 2.2 | 1 | 100.00 |
| *Empedobacter* | 1 | 2.2 | 0 | 0.00 |
| *Enterococcus* | 1 | 2.2 | 0 | 0.00 |
| *Eschericia* | 1 | 2.2 | 1 | 100.00 |
| *Klebsiella* | 4 | 8.7 | 1 | 25.00 |
| *Ledercia* | 1 | 2.2 | 1 | 100.00 |
| *Pantoea* | 2 | 4.3 | 2 | 100.00 |
| *Providencia* | 1 | 2.2 | 0 | 0.00 |
| *Pseudomonas* | 1 | 2.2 | 0 | 0.00 |
| *Stenethophomonas* | 1 | 2.2 | 0 | 0.00 |
| *Waustersiella* | 1 | 4.3 | 0 | 0.00 |
| **Total** | **46** | **100** | **17** | **36.96** |

**Table 4.** Summary of the resistance patterns of the isolates obtained (by species).

| ISOLATE | Gram<br>reaction | Resistance |  | Total (%) |
| --- | --- | --- | --- | --- |
|  |  | No | Yes |  |
| *Acinetobacter baumannii* | Negative | 0 | 2 | 2 (4.35) |
| *Acinetobacter haemolyticus* | Negative | 4 | 1 | 5 (10.87) |
| *Acinetobacter junii* | Negative | 2 | 0 | 2 (4.35) |
| *Acinetobacter lwoffii* | Negative | 3 | 0 | 3 (6.52) |
| *Acinetobacter sps\** | Negative | 3 | 3 | 6 (13.04) |
| *Acinetobacter variabilis* | Negative | 0 | 1 | 1 (2.17) |
| *Citrobacter freundii* | Negative | 0 | 1 | 1 (2.17) |
| *Empedobacter brevis* | Negative | 1 | 0 | 1 (2.17) |
| *Enterobacter cloacae* | Negative | 1 | 4 | 5 (10.87) |
| *Enterococcus faecium* | Negative | 1 | 0 | 1 (2.17) |
| *Eschericia coli* | Negative | 0 | 1 | 1 (2.17) |
| *Klebsiella oxytoca* | Negative | 1 | 0 | 1 (2.17) |
| *Klebsiella pneumoniae* | Negative | 1 | 1 | 2 (4.35) |
| *Ledercia adecarboxylata* | Negative | 0 | 1 | 1 (2.17) |
| *Pantoea calida* | Negative | 0 | 1 | 1 (2.17) |
| *Pantoea dispersa* | Negative | 0 | 1 | 1 (2.17) |
| *Pseudomonas stutzeri* | Negative | 1 | 0 | 1 (2.17) |
| *Staphylococcus aureus* | Positive | 6 | 0 | 6 (13.04) |
| *Stenethophomonas maltophilia* | Negative | 1 | 0 | 1 (2.17) |
| *Klebsiella varicola* | Negative | 1 | 0 | 1 (2.17) |
| *Providencia rettgeri* | Negative | 1 | 0 | 1 (2.17) |
| *Waustersiella falseniia* | Negative | 2 | 0 | 1 (4.35) |
| **Total (%)** | **Negative-**<br>**86.96%**<br>**Positive-**<br>**13.04%** | **29**<br>**(63.04)** | **17**<br>**(36.96)** | **46 (100)** |

\*This organism was only identified to genus level

The resistance markers tested included ESBL, carbapenem resistance (CRO), methicillin sensitivity for *Staphylococcus aureus*, and vancomycin sensitivity. Of all the 46 isolates obtained, significant resistance was observed. 10.9% (n=5) showed both ESBL and CRO resistance, whole 23.9%(n=11) were positive for ESBL production. All the rest were non-resistant strains as shown by negative ESBL at 47.8%(n=22), methicillin sensitive at 13% (n=6) and vancomycin sensitivity at 2.2%(n=1) **(Table 5)**.

**Table 5.**
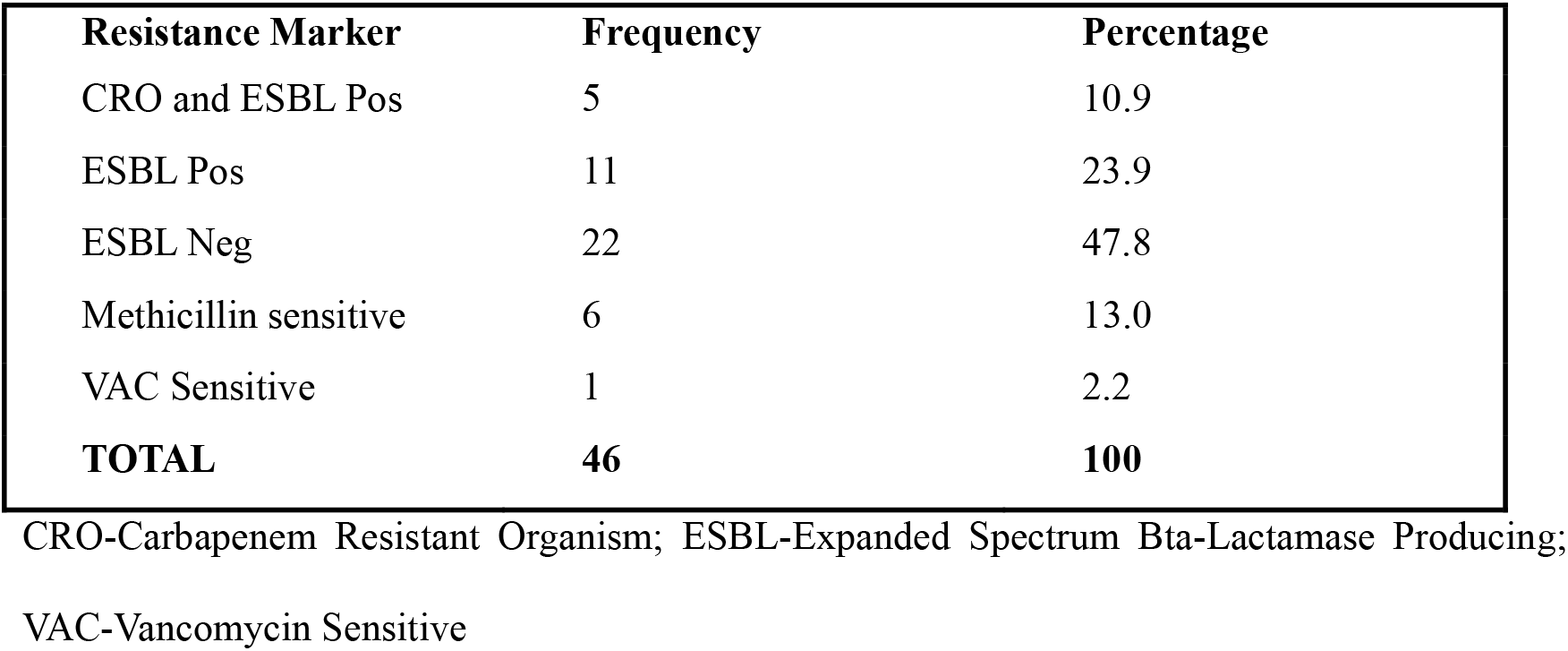
Summary of resistance makers

| **Resistance Marker** | **Frequency** | **Percentage** |
| --- | --- | --- |
| CRO and ESBL Pos | 5 | 10.9 |
| ESBL Pos | 11 | 23.9 |
| ESBL Neg | 22 | 47.8 |
| Methicillin sensitive | 6 | 13.0 |
| VAC Sensitive | 1 | 2.2 |
| **TOTAL** | **46** | **100** |

CRO-Carbapenem Resistant Organism; ESBL-Expanded Spectrum Bta-Lactamase Producing; VAC-Vancomycin Sensitive

## DISCUSSION

A swab positivity of 61.3% among the surfaces was considered high and indicative of a high level of contamination across the departments. Previous studies have found varying contamination rates among frequented surfaces, equipment, and Health Care Workers’ (HCW\0’ hands. In one such study, a contamination rate of 85.8% was reported on stethoscopes used at a tertiary hospital(13). In a similar setting in Tanzania, a contamination rate of 26.4% was noted among HCWs’ hands(10). In similar hospitals in Kenya, diverse but close contamination rates have also been reported, including 55% by Nyaruaba et al. (14) and 95.9% by Odoyo et al. (15). In the study by Odoyo et al. (15), potential factors that could lower this high contamination rate were also studied. The study recommended daily washing of patient bedding, provision, and education on the use of handwashing stations, supply of running water for handwashing, and soap for handwashing lowered the bacterial load of the considered surfaces significantly(15).

Hence, based on these recommendations, there is a need to perform a study that assesses adherence to IPC guidelines. Since Kenya introduced IPC guidelines in 2019(16) to reduce HAIs, numerous challenges have limited their effective implementation, including resource limitations such as lack of personal protective equipment for HCWs, lack of running water, and general poor adherence to hygiene regulations(17).

The contaminants reported were mainly gram-negative bacteria, with *Staphylococcus spp* being the only gram-positive organism. This was similar to the isolates reported in similar studies in Kenya(14,15) and Tanzania(10). In the study by Chaoui et al. (11) in Morocco, the primary isolates obtained on the surfaces swabbed were *Enterobacteria* (31.6%), *Staphylococcus aureus* (24%), *Pseudomonas aeruginosa* (9.2%) and *Acinetobacter spp*. (3.3%). It is worth noting that contrary to the Kenyan studies, where *Acinetobacter* or *Escherichia*(8,9,14,15,18) are the most commonly observed bacterial contaminants, the North African study had *Acinetobacter* as the least prevalent and *Enterobacteria* as the most common(11). Potential explanations for this difference have not been presented in literature. Overall, the most common contaminants in hospital environments are Gram-negative bacteria and *Staphylococcus spp*. (3,4,17). Is it probable that a country’s development status would influence the prevalence and types of contaminants reported in the hospital environment? This remains to be answered.

Raoofi et al.(19), in a meta-analysis that pooled the global prevalence of HAIs, reported that HAI rates were higher in Africa and the developing world and generally lower in America, Europe, and developed countries. Unfortunately, the differences in the types of bacteria causing HAIs in these regions are yet to be extensively studied and summarized. This may be attributed to resource availability, well-designed healthcare systems, and adherence to IPC guidelines in the developed countries(19). In a systematic review that pooled data mainly from developed countries in Europe and America, Saleem et al. (20) concluded that *Klebsiella pneumonia, Pseudomonas aeruginosa* and *E. coli* were the most reported pathogens. Still, gram negative pathogens were most common causes of HAIs, similar to data reported in Africa(10,21) and even Kenya(8,14,15). Since evidence has been presented that high rate of contamination translates to high HAI rates(22), the need to invest more in hospital environment decontamination has never been more urgent(23).

In a discussion of the central role of hospital environmental contamination in favoring the spread of multi-drug resistant Gram Negative pathogens, Chia et al. (12) posit that there is increasing evidence to support the hypothesis that hospital contamination with resistant pathogens may directly translate to higher prevalence of highly resistant HAIs. The fact that the most common contaminants reported across the board are members of the stubborn ESKAPE group supports this belief(9). For instance, in a similar study among five Kenyan hospitals, Odoyo et al. (9) observed that all the hospital departments sampled were contaminated with MDR ESKAPE organisms. 95.6% of the isolates of *A. baumannii, Enterobacter* species, and *K. pneumoniae* were resistant to common antibiotics like piperacillin, ceftriaxone, and cefepime. More alarming was that all the *K. pneumoniae* isolates were resistant to all antibiotics tested except colistin(9). These findings, which have been confirmed in our study, were similar to those reported in other studies(14,15,18).

Significant swab positivity was observed in all the departments sampled, with the gynecology, pediatric, and newborn unit having more special positivity rates similar to the findings of Odoyo et al. (9). Among developing countries, surgical site infections have been reported as the most common HAIs(27). Thus, with caesarian sections being the most common surgeries in Kenya(28), the risk to the mother and the neonate is always more elevated. The high contamination rate reported in this study, because the hospital environment poses a transmission risk to patients, may partly explain the high rate of HAIs reported in developing countries(9,29).

The fact that resistant pathogens were reported across all the departments is also worth considering in this interpretation. MDR pathogens, especially those of the ESKAPE family, initiate more severe HAIs, which may badly affect patients in these departments(30). Since resistant pathogens were isolated in the NBU and pediatric wards as well, the risk they pose to the immune-naïve populations in these departments is worth contemplating. Newborns would generally be more affected by HAIs since their immune system is underdeveloped (31). The fact that resistant pathogens are surface contaminants in departments with some of the most vulnerable patients in the hospital should point to an urgent need to adhere strictly to IPC guidelines to mitigate the situation(7,31,32). Whereas the present study did not determine the statistical association between the department from which a specimen was obtained and the occurrence of resistance phenotypes, the data showed that the paediatric and NBU units were most affected by these. This aspect of association has also yet to be considered by any other studies, hence a need to statistically quantify this association through inferential statistics.

Generally, some bacterial species have a documented higher rate of AMR than others. The members of the ESKAPE family are a case in point(6). In the present study, *Acinetobacter spp* were the most reported contaminants and doubled up as the genus with the highest resistance rate. A similar finding had also been reported in an analysis that considered five different hospitals in Kenya(9). The increased contamination rate with *Acinetobacter spp* is noteworthy owing to the bacteria’s unique characteristics that give it the ability to cause HAIs. First is its ability to form biofilms which has been confirmed by Espinal and Vila (33). Additionally, its ability to survive through long periods of desiccation (34). These properties make *Acinetobacter spp* an important clinical pathogen implicated in many HAIs, including ventilator-associated pneumonia, skin and soft tissue infections, urinary tract infections, secondary meningitis, and bloodstream infections in Kenya as other parts of the globe(9,35,36). The pathogen has been implicated in several HAI reports in Kenya(37) and one case, as a cause of a hospital outbreak(38). The fact that *A. boumanii* is intrinsically resistant to many of the commonly used antibiotics makes it a time bomb that is waiting to explode, hence the need to give it immediate attention(9,15,39). Thus, there is a pressing need to direct more effort toward hospital environmental hygiene to reduce contaminants and prevent the possibility of HAI outbreaks, especially those caused by resistant pathogens.

Other gram-negative pathogens observed in this study, in addition to the gram positive *Staphylococcus aureus* are not to be given a blind eye in the implementation of hospital IPC strategies. According to Chia et al. (12), MDR gram negative organisms are associated with high mortality rate and continue to present a challenge to healthcare systems globally. Since there is a growing body of knowledge to associate hospital environmental contamination and occurrence of HAIs(22), hospital IPC guidelines should be made as comprehensive as possible. In support of this, Chaoui et al. (11) detailed that the idea of environmental bacterial reservoir is a reality that poses a public health risk hence needing strict current recommendations for hand hygiene, cleaning, and disinfection of surfaces in hospitals. In a study by Darge et al.(21) 68.4% gram positive and 31.5% gram negative isolates were identified as contaminants in a hospital in Ethiopia, with both category showing significance AMR. Moremi et al. (40) argued that room occupancy by a patient shedding nosocomial pathogens may the primary source of surface contaminants, thus drawing attention to special patient considerations as part of IPC guidelines. This finding also agreed with conclusions from a meta-analysis by Mitchel et al. (41). It is also possible that general surface contamination may also be a source of these pathogens. As explained by Hajar et al. (42), dispersion of gram negative contaminants from colonized sinks to hands of staff may also happen when these sinks are not thoroughly cleaned. Sinks are among the frequented surfaces in the hospital setting, together with others like shared equipment and door handles. To combat the high contamination rate, there is a need to improve overall adherence to IPC guidelines, with more emphasis being placed on clearing surface contamination.

## CONCLUSION

A high contamination rate with pathogenic bacterial species was noted among the departments sampled. Among the isolates obtained, also, a significant rate of antibiotic resistance was observed. *Acinetobacter spp* was the most common pathogen isolated and doubled up as the one with the highest resistance to common antibiotics. We, therefore, recommend thorough adherence to IPC guidelines and regular surveillance as the best approaches to mitigate the problem. Genomic-level studies also need to map the transmission of resistant pathogens between the environment and patients.

## DECLARATIONS

### Ethical Approval

The study was approved by the KEMRI Scientific Ethics Review Unit and assigned approval number SERU/CGMR-C/054/3318. Further approval before data collection was also obtained from the facility management. Staff whose hands were swabbed gave consent for the procedure.

### Consent for publication

Not applicable

### Availability of data and materials

Not applicable

### Authors’ contributions

The study was conceptualized by Kolek Chester and compiled by Kavulavu Briton. It was then reviewed by Faith Okalebo, Benson Singa and Kirkby D Tickell. Actual data collection was completed by Mary Masheti and Chris Oduol. Ian Omuom, Ochieng Odhoch, being part of the hospital leadership, reviewed the study and assisted in data collection. Actual laboratory work was done by Caroline Tigoi and Robert Musyimi. Data analysis was completed by Kolek Chester, Kavulavu Briton, and Faith Okalebo who also produced the tables and figures. The manuscript was developed by Kavulavu Briton and Kolek Chester and reviewed by all co-autors before submission.

## Acknowledgements

The authors wish to specifically thank the management and staff of Migori County Referral Hospital for facilitating the data collection process.

